# A kinase-dead *Csf1r* mutation associated with adult-onset leukoencephalopathy has a dominant-negative impact on CSF1R signaling

**DOI:** 10.1101/2021.09.29.462493

**Authors:** Jennifer Stables, Emma K. Green, Anuj Sehgal, Omkar Patkar, Sahar Keshvari, Isis Taylor, Maisie E. Ashcroft, Kathleen Grabert, Evi Wollscheid-Lengeling, Stefan Szymkowiak, Barry W. McColl, Antony Adamson, Neil E. Humphreys, Werner Mueller, Hana Starobova, Irina Vetter, Sepideh Kiani Shabestari, Matthew M. Blurton-Jones, Kim M. Summers, Katharine M. Irvine, Clare Pridans, David A. Hume

**Affiliations:** Mater Research Institute-University of Queensland, Translational Research Institute, Brisbane, Qld, Australia; Centre for Inflammation Research and Simons Initiative for the Developing Brain, University of Edinburgh, Edinburgh, UK; Toxicology Unit, Institute of Environmental Medicine, Karolinska Institutet, Stockholm, Sweden; Luxembourg Centre for Systems Biomedicine. Université du Luxembourg, Belvaux, Luxembourg; UK Dementia Research Institute, Centre for Discovery Brain Sciences, University of Edinburgh, Edinburgh, UK; Genome Editing Unit, Faculty of Biology, Medicine and Health, University of Manchester, Manchester, UK; Institute for Molecular Biosciences & School of Pharmacy, University of Queensland, Brisbane, Australia; Department of Neurobiology & Behavior, University of California, Irvine CA. USA

**Keywords:** CSF1R, macrophage, kinase-dead, leukoencephalopathy

## Abstract

Amino acid substitutions in the kinase domain of the human *CSF1R* gene are associated with autosomal dominant adult-onset leukoencephalopathy with axonal spheroids and pigmented glia (ALSP). To model the human disease, we created a disease-associated mutation (Glu631Lys; E631K) in the mouse *Csf1r* locus. Homozygous mutation (*Csf1r*^E631K/E631K^) phenocopied the *Csf1r* knockout; with prenatal mortality or severe postnatal growth retardation and hydrocephalus. Heterozygous mutation delayed the postnatal expansion of tissue macrophage populations in most organs. Bone marrow cells from *Csf1r*^E631K/+^ mice were resistant to CSF1 stimulation *in vitro*, and *Csf1r*^E631K/+^ mice were unresponsive to administration of a CSF1-Fc fusion protein which expands tissue macrophage populations in controls. In the brain, microglial cell numbers and dendritic arborization were reduced in the *Csf1r*^E631K/+^ mice as in ALSP patients. The microglial phenotype is the opposite of microgliosis observed in *Csf1r*^+/-^ mice. However, we found no evidence of brain pathology or impacts on motor function in aged *Csf1r*^E631K/+^ mice. We conclude that disease-associated *CSF1R* mutations encode dominant negative repressors of CSF1R signaling. We speculate that leukoencephalopathy associated with human CSF1R mutations requires an environmental trigger and/or epistatic interaction with common neurodegenerative disease-associated alleles.

**Summary Statement:** This study describes the effect of a human disease-associated mutation in the mouse CSF1R gene on postnatal development and growth factor responsiveness of cells of the macrophage lineage.

## Introduction

The colony stimulating factor 1 receptor gene (*Csf1r*) encodes a ligand-dependent tyrosine kinase receptor that controls the survival, proliferation, and differentiation of mononuclear phagocyte populations throughout the body including microglia in the brain (Chitu and Stanley, 2017; Stanley and Chitu, 2014). CSF1R has two ligands, colony stimulating factor 1 (CSF1) and interleukin 34 (IL-34) (Lelios et al., 2020). Upon ligand binding, CSF1R dimerization and autophosphorylation generates phosphotyrosine motifs that act as docking sites for multiple downstream effector pathways (Chitu and Stanley, 2017; Stanley and Chitu, 2014). Biallelic recessive loss-of-function mutations in mouse, rat and human *CSF1R* genes are causally linked to osteopetrosis and postnatal developmental abnormalities (reviewed in (Chitu et al., 2021; Hume et al., 2020)). In 2011, Rademakers *et al* (Rademakers et al., 2011) reported heterozygous amino acid substitutions in the tyrosine kinase domain of CSF1R in patients with autosomal dominant adult-onset leukoencephalopathy with axonal spheroids and pigmented glia (ALSP), now also called CSF1R-related leukoencephalopathy (CRL) (Chitu et al., 2021). Since that time, more than 100 different disease-associated *CSF1R* coding mutations have been identified (Chitu et al., 2021; Guo and Ikegawa, 2021; Konno et al., 2018; Konno et al., 2017). Characteristic features of ALSP include enlarged ventricles, cerebral atrophy, periventricular calcifications and thinning of the corpus callosum (Konno et al., 2018; Konno et al., 2017). In ALSP brain the microglia are reduced in number and altered in their morphology and gene expression (Kempthorne et al., 2020; Tada et al., 2016).

To understand the molecular basis of ALSP, we transfected factor-dependent Ba/F3 cells with expression vectors encoding wild-type CSF1R or disease-associated mutant receptors (Pridans et al., 2013). The mutant CSF1R proteins were expressed on the cell membrane at similar levels to wild-type CSF1R and bound and internalised CSF1, but were unable to support CSF1-dependent cell survival or proliferation. These findings support a dominant-negative model for ALSP. In patients, inactive homodimers and heterodimers may also compete for ligands with the functional receptor dimers (Hume et al., 2020). Other authors have suggested, based upon analysis of a heterozygous *Csf1r* knockout (*Csf1r*^+/-^) in C57BL/6J mice that the dominant inheritance in ALSP arises from CSF1R haploinsufficiency (Arreola et al., 2021; Biundo et al., 2020; Chitu et al., 2020; Chitu et al., 2015). However, these models are associated with microgliosis. Neither the microgliosis phenotype, nor changes in microglia-specific gene expression, was replicated in *Csf1r*^+/-^ rats (Patkar et al., 2021a). Similarly, there is no reported evidence of neuropathology in aged obligate carriers of recessive *CSF1R* loss-of-function alleles in humans (Guo et al., 2019; Guo and Ikegawa, 2021).

Microglia have been ascribed numerous roles in brain development and maturation (Prinz et al., 2019). Against that background, the generation of a hypomorphic *Csf1r* mutation in mice that was entirely microglia-deficient but developed normally was surprising (Rojo et al., 2019). The *Csf1r*^ΔFIRE^ mutation removed a conserved intronic enhancer required for expression of the receptor in microglia. *Csf1r*^ΔFIRE/ΔFIRE^ mice also lacked CSF1R expression in bone marrow (BM) progenitors and blood monocytes and were deficient in resident macrophage populations in skin, peritoneum, kidney and heart (Rojo et al., 2019). These observations indicate that microglia and certain peripheral macrophage populations are uniquely dependent upon CSF1R and/or have distinct transcriptional regulation.

Konno *et al*. (Konno et al., 2014) reported that transient over-expression of kinase-dead mutant CSF1R proteins in HEK293 cells stably expressing wild-type (WT) CSF1R did not inhibit CSF1-induced autophosphorylation. This finding has been cited as evidence against a dominant-negative model (Chitu et al., 2021). Here we describe the generation and characterisation of mice carrying the disease-associated *Csf1r* E631K mutation (Glu633Lys in human), one of the kinase-dead mutations analysed previously in Ba/F3 cells (Pridans et al., 2013). The results confirm the dominant-negative effect on CSF1R signalling and provide novel insights into macrophage homeostasis and the limitations of mice as models of microglial homeostasis.

## Materials and Methods

### CRISPR/Cas9 design

The methods used for targeted mutagenesis are similar to those previously used to insert a Fusion Red cassette into the mouse *Csf1r* locus (Grabert et al., 2020). Guides were designed using the Sanger website with stringent criteria for off target predictions (guides with mismatch (MM) of 1 or 2 for elsewhere in the genome were discounted). Single stranded RNA (ssRNA) guide 513 (g513) was used to cut the sequence around the glutamate codon at position 631 of *Csf1r* on chromosome 18. A ssDNA template then induced the base change GaG to AaA converting to a lysine codon and deleted the Alul site to allow for genotyping (**Fig. S1A**). An Alt-R crRNA (IDT) oligo was resuspended in sterile RNase free injection buffer (Tris HCl 1 mM, pH 7.5, EDTA 0.1 mM) and annealed with transactivating crispr RNA (tracrRNA);IDT) by combining 2.5 μg crRNA with 5 μg tracrRNA and heating to 95 °C. The mix was left to slowly cool to room temperature (RT). After annealing the complex an equimolar amount was mixed with 1000 ng Cas9 recombinant protein (NEB; final concentration 20 ng/μL) and incubated at RT for 15 min, before adding Cas9 messenger RNA (mRNA) (final concentration; 20 ng/μL) and the ssDNA polyacrylamide gel electrophoresis (PAGE) purified repair template (IDT; final concentration 50 ng/μL) in a total injection buffer volume of 50 μL. The injection mix was centrifuged for 10 min at RT and the top 40 μL removed for microinjection.

### Generation of C57BL/6J.Csf1r^Em1Uman^ (Tg16) mice

Microinjections were performed using AltR crRNA:tracrRNA:Cas9 complex (20 ng/μL; 20 ng/μL; 20 ng/μL respectively), Cas9 mRNA (20 ng/μL), and ssDNA homology-directed repair (HDR) template (50 ng/μL). The mix was injected into one-day single cell mouse embryos (C57BL/6JOlaHsd). The zygotes were cultured overnight, and the resulting 2 cell embryos implanted into the oviduct of day 0.5 post-coitum pseudopregnant mice.

### Genotyping

Genotyping was performed using the Phire Direct Tissue PCR kit using the Storage and Dilution protocol recommended by the manufacturer. In brief, ear clips were digested then PCR performed at an annealing temperature of 62 °C with forward (5’ACGCCTGCATTTCTCATTCC) and reverse primers (5’ATCCAGCTCTTACCTCCGTG).

The DNA was then digested with 2 μL CutSmart Buffer (NEB, R0137L) and 1 μL Alul enzyme (NEB, R0137L) at 37 °C for 1 h. The products were then separated on a 2% agarose gel. The expected products were 121 bp, 77, bp, 6 bp (+/+), 198 bp, 121 bp, 77 bp, 6bp (+/E631K) and 198 bp, 6 bp (E631K/E631K) as per **Fig. S1B**. For ongoing breeding a qPCR-based protocol was also devised. Two qPCR reactions were run in parallel, using a forward primer specific for wild type *Csf1r* (WT-F: AAGGAGGCCCTGATGTCAGAG) or *Csf1r-*E631K (MUT-F: AAGGAGGCCCTGATGTCAAAA) with a universal reverse primer (R: ACAGGCTCCCAAGAGGTTGA). The wild type *Csf1r* allele was poorly amplified using the mutant primer and vice versa (delta Ct ~10). *Csf1r-*EGFP mice were genotyped by qPCR using GFP-specific primers (F: ACTACAACAGCCACAACGTCTATATCA, R: GGCGGATCTTGAAGTTCACC).

### Animal breeding

C57BL/6J.Csf1r^Em1Uman^ (Tg16) donor and recipient mice were C57BL/6JOlaHsd. They were then crossbred on a C57BL/6JCrl background with interbreeding of the offspring then further backcrossed to C57BL/6JCrl. Subsequent to the transfer to Australia, the mice were rederived and bred and maintained in specific pathogen free facilities at the University of Queensland (UQ) facility within Translational Research Institute. To enable visualisation of myeloid populations in tissues, the *Csf1r*^E631K^ line was bred to the *Csf1r*-EGFP reporter transgenic line (Sasmono et al., 2003) also backcrossed >10 times to the C57BL/6JArc genetic background. For comparative analysis, mice bearing the *Csf1r*^ΔFIRE^ hypomorphic allele (Rojo et al., 2019) were transferred from Edinburgh to UC Irvine and bred to the C57BL/6J genetic background. For analysis of embryos, timed matings were set up in the late afternoon. If a plug was detected next morning this was considered as embryonic day (E) 0.5 (E0.5).

### Animal Ethics

In the UK, ethical approval was obtained from The University of Edinburgh’s Protocols and Ethics Committees under the authority of a UK Home Office Project License under the regulations of the Animals (Scientific Procedures) Act 1986. In Australia, all studies were approved by the Animal Ethics Committee of the University of Queensland. Mice were housed and bred under specific pathogen free conditions.

### Tissue processing

Embryos were fixed in 10% neutral buffered formalin for 2 days and then transferred to 70% ethanol. Embryos were cut in half along the sagittal plane. For postnatal analyses, peripheral blood and peritoneal cells were collected as previously described (Rojo et al., 2019). Following peritoneal lavage, tissues of interest were removed. Tissue for qPCR analysis was snap frozen in TRI Reagent. Tissues for immunohistochemistry (IHC) were post-fixed in 4% PFA for ~6 hours, then transferred to 1x Phosphate Buffered Saline (PBS) PBS with 0.01% sodium azide. Tissues were embedded in paraffin using standard methods by core histology facilities at the Queen’s Medical Research Institute, Edinburgh or the Translational Research Institute, Brisbane.

### Immunohistochemistry

Sources of antibodies and other reagents are provided in **Table S1**.

For IHC of embryos, antigen retrieval was performed with Vector Antigen Unmasking solution at 100 °C for 5 min. Non-specific protein binding was blocked with 2.5% horse serum for 20 min. For microglia detection, slides were incubated with rabbit anti-IBA1 primary antibody (1:2000) at room temperature for 30 min. After washing in phosphate-buffered saline (PBS), slides were incubated with secondary antibody at RT for 35 min. Following two washes in PBS, slides were incubated with peroxidase substrate for 5 min. Slides were finally washed and counterstained with haematoxylin, then dehydrated before mounting with Pertex mounting medium.

Tissues for whole mount imaging were extracted and kept in PBS on ice until imaged with an Olympus FV3000 confocal microscope. For IHC analysis of adults, tissues were fixed and processed for paraffin-embedded histology using routine methods, with the exception of 7-week-old brains which were reserved for free-floating IHC. These brains were cryoprotected, frozen in OCT and 40um serial, coronal sections were collected in a rostro-caudal manner (1 in 12 series) using a Leica CM1950 cryostat. 7-week-old spleens were fixed, cryoprotected, and frozen in OCT.

Frozen spleen sections (5 μm) were sequentially stained with F4/80 and CD169. Free-floating brain sections were incubated at RT for 30 min in permeabilization buffer (0.1% Triton-X in PBS) followed by 90 min in blocking solution (5% NGS, 0.3% Triton-X in PBS). Sections were then incubated for overnight at 4° C under orbital agitation in the primary antibody against defined surface markers. Following 3 × 10 min washes in permeabilization buffer, slices were incubated in the appropriate secondary antibody (**Table S1**) diluted in blocking solution, for 90 min at RT in the dark. Slices were then washed in permeabilization buffer, followed by a 5 min incubation with 4′,6-diamidino-2-phenylindole (DAPI) diluted in PBS, and a further 10 min wash in permeabilization buffer. All sections were washed with PBS for 5 min and mounted with Fluorescence Mounting Medium. Images were acquired on an Olympus FV3000 confocal microscope.

GFAP^+^, P2RY12^+^, and TMEM119^+^-areas were quantified using ImageJ (https://imagej.net/). The percentage area of positive staining was calculated using the ‘measure’ tool in ImageJ, following adjustment of brain region specific threshold, which was kept consistent for all mice. To calculate IBA1+ cell body size, maximum intensity projections were opened in Fiji v1.5 and the free drawing tool was used to measure the area of positive cells with a visible nucleus (20 cells per animal). For microglial arborisation, images were analysed as previously described (Patkar et al., 2021b).

Paraffin-embedded tissues were sectioned at 6 μm using a Leica RM2245 microtome. F4/80 and IBA1 staining was performed as previously described (Keshvari et al., 2021). Whole-slide digital imaging was performed on the VS120 Olympus slide scanner. IBA1+ density was analysed in four different fields per sample. For liver F4/80 quantification, the DAB-positive areas were quantified as a percentage using the ‘measure’ tool in ImageJ, following adjustment of the image threshold, which was kept consistent for all mice. To quantify the corpus callosum area identified by staining with Luxol fast blue (Pridans et al., 2018) the free drawing tool in Fiji was used to measure the area of the cerebral hemisphere, and then the corpus callosum contained within that hemisphere. The area of the corpus callosum as a % of the total area of the cerebral hemisphere was then calculated.

### Embryo image acquisition and quantification

Images were acquired using the NanoZoomer (Hamamatsu) slide scanner at 40X magnification. Image analysis was performed with NDP.view software (Hamamatsu) and ImageJ. For the embryo analysis, images were exported as tiff format from the NDP.view files. In ImageJ, the perimeter of the developing brain was traced, and the area measured. Colour threshold settings were used to remove the white gaps in the developing brain for an exact area of brain. Further colour threshold settings were applied to measure the IBA1 staining and then a percentage of IBA1 staining in the whole developing brain was calculated per embryo. For quantification of IBA1 staining in the embryo livers the same ImageJ protocol was followed.

### Analysis of response to CSF1 in vitro and in vivo

Bone marrow cells were harvested and cultured in varying concentrations of recombinant human CSF1 as described previously (Rojo et al., 2019). After 7 days, 25ug/mL of resazurin was added to each well. Plates were returned to the incubator for 1 hour. Optical density was then measured on a plate reader (Omega Pherastar, BMG Labtech).

To assess *in vivo* response to CSF1, mice were injected with a recombinant human-CSF1 mouse Fc conjugate (Novartis, Switzerland). 6-week-old littermates received one injection per day of 5mg/kg of CSF1-Fc for 4 days between Zeitgeber time (ZT) 2-3.

### Micro-CT imaging and reconstruction

Left-hind limbs (LHL) were fixed in 4% paraformaldehyde (PFA) and subsequently transferred into PBS for high-resolution micro-computed tomography (μCT) scanning using Bruker’s Skyscan 1272 (Bruker, BE). X-ray settings were standardised to 70 kV and 142 μA. The x-ray filter used was a 0.5 mm aluminium. The entire femora were scanned over 360° rotation in 0.8° rotational steps and the exposure time was set to 470 ms. Projections were acquired with nominal resolutions of 10 μm and each slice contained 1224 × 820 pixels. All X-ray projections were reconstructed using a modified back-projection reconstruction algorithm (NRecon 1.7.3.1 software-SkyScan, Bruker) to create cross-sectional images. Reconstruction parameters included ring artefact correction (2–6), beam hardening correction (40–50%), and misalignment correction. Reconstruction was performed in a blinded manner. 3D reconstructions were viewed using CTvox 3.3.0 (Bruker). Reconstructed images were analyzed through CTAn 1.19 software (Bruker) which has inherent 2D and 3D analysis tools. All analysis was performed as per the updated guidelines for the assessment of bone density and microarchitecture *in vivo* using high-resolution peripheral quantitative computed tomography (Whittier et al., 2020).

### Magnetic resonance imaging

MRI was performed on a 7 T horizontal bore Biospec AVANCE neo preclinical imaging system equipped with a 116 mm bore gradient insert (Bruker BioSpin GmbH, Germany, maximum gradient strength 660 mT/m). Mice were anaesthetised with 1.5–2% isoflurane (Zoetis Ltd., London UK) in oxygen/air (50/50, 1□L/min) and secured in a cradle (Rapid Biomedical GmbH, Rimpar, Germany). The respiration rate and rectal temperature were monitored (Model 1030 monitoring and gating system, Small Animal Instruments Inc. Stony Brook, NY, USA), with body temperature maintained at 37°C by a heat fan. An 86 mm quadrature volume coil (Bruker BioSpin GmbH, Germany) was used for transmission with signal reception by a two-channel phased-array mouse brain coil (Rapid Biomedical GmbH, Rimpar, Germany).

Scout images were taken to confirm correct positioning and the magnetic field was optimised using automated 3D field mapping routine. For all subsequent sequences, the field of view was 19.2 × 19.2 mm and the slice thickness 0.8 mm. For anatomical imaging, 17 coronal slices covering the entire brain were acquired using a T2-weighted Rapid Acquisition with Relaxation Enhancement (RARE) sequence with the following parameters: matrix size 192 × 192, TR 2300 ms, effective TE 36 ms, RARE factor 4, number of signal averages 4. The scan time was 7 min 21 s.

### Flow cytometry

Microglial cells were isolated as described by Grabert et al. (2020). Myeloid lineage, HSPC and committed progenitor subsets phenotyping was performed on BM suspensions as detailed in **Table S1.** Peripheral blood, peritoneal lavage cells and isolated microglia were also stained for myeloid lineages as detailed in **Table S1.** Cell acquisition was performed on Beckman Coulter’s Cytoflex Analyser (Beckman Coulter, USA) or BD LSRFortessa™ X-20 (BD Biosciences, USA). Data analysis was performed using the FlowJo software (Tree Star Data Analysis Software, Ashland, OR).

### IGF1 and CSF1 Immunoassay

Serum IGF1 and serum CSF1 were measured using commercial kits (**Table S1**) according to manufacturer’s instructions.

### RNA purification and qRT-PCR analysis

mRNA isolation, quantification and assessment of integrity were carried out as described previously (Keshvari et al., 2021) and gene expression was quantified using the SYBR Select Master Mix on an Applied Biosystems QuantStudio real-time PCR system. Gene expression relative to *Tata* was calculated using the delta Ct method.

Primer sequences used were:

Mmp9 – F: AGGGGCGTGTCTGGAGATTC, R: TCCAGGGCACACCAGAGAAC
Plau – F: GGTTCGCAGCCATCTACCAG, R: TTCCTTCTTTGGGAGTTGAATGAA
Tata – F: CTCAGTTACAGGTGGCAGCA, R: ACCAACAATCACCAACAGCA

### Sensorimotor testing

Sensorimotor testing was conducted on 43-week-old mice. In all tests, the experimenter was blinded as to the genotype of each mouse. Mechanical paw withdrawal threshold was measured using an electronic von Frey apparatus (MouseMet Electronic von Frey, Topcat Metrology Ltd, Little Downham, United Kingdom). Mice were habituated in individual mouse runs for 30 minutes prior to commencing measurement. As described previously (Hasan et al., 2021) a soft-tipped von Frey filament, was placed against the foot pad of the right hind paw. Pressure was slowly increased at a rate of 1 g/s, through rotation of the device handle and the force (g) causing paw withdrawal displayed on the device was recorded. A single biological replicate was determined by averaging three repeated measurements (minimum of 5 min intervals) for each mouse (Hasan et al., 2021).

Locomotor performance was measured using the Parallel Rod Floor apparatus (Stoelting Co, Wood Dale, IL). Each mouse was placed into the centre of the apparatus. The total distance travelled (m) and number of errors (foot slips off the rods) were recorded for over a period of 2 mins and analysed using the ANY-Maze software (Stoelting Co.). Gait analysis was performed using the CatwalkXT (Noldus Information Technology, Wageningen, Netherlands). Animals were allowed to voluntarily traverse the enclosed, illuminated glass surface. All recordings were performed in a dark room. A camera captured the illuminated footprints from below the glass surface to record the paw placement of each mouse as it traversed the platform. Only runs of 3- to 12-second duration with speed variances below 90% were considered acceptable. The mouse remained on the platform until 3 acceptable runs had been recorded. Analysis of these runs was performed using the CatwalkXT software.

## Results

### Generation of C57BL/6J.Csf1r^Em1Uman^ (Tg16) mice

To create C57BL/6J.Csf1r^Em1Uman^ (Tg16) mice, gRNAs of tracrRNA, crispRNA and Cas9 were microinjected as described previously (Grabert et al., 2020). Nine founder mice (*Csf1r*^+/E631K^) were crossed with C57BL/6JCrl mice and the offspring interbred. In the initial analysis undertaken in Edinburgh, the frequencies of *Csf1r*^E631K/E631K^ embryos up to E14.5 were not significantly divergent from Mendelian expectation, but no live homozygous pups were obtained. *Csf1r*^E631K/E631K^ embryos were found previously to be almost entirely macrophage and microglia-deficient when analysed as controls for the more selective loss of microglia in *Csf1r*^ΔFIRE/ΔFIRE^ mice (Munro et al., 2020). Subsequent to transfer of the mutant line to Australia, and cross to the C57BL/6JArc background, the *Csf1r*^E631K^ allele was crossed to the *Csf1r*-EGFP reporter transgenic line (Sasmono et al., 2003) on the same genetic background to enable visualisation of tissue macrophage populations by whole mount imaging.

*Csf1r* mRNA is expressed in the earliest macrophages produced in the yolk sac (Lichanska et al., 1999) and subsequent development of macrophages in the mouse embryo depends upon CSF1R signalling (Hoeffel et al., 2015; Munro et al., 2020). **Figure 1** shows representative histology and the localisation of embryonic macrophages expressing IBA1 in *Csf1r^+/+^, Csf1r*^E631K/+^ and *Csf1r*^E631K/E631K^ embryos at E12.5. The three genotypes were indistinguishable at this age (**Figure 1A**). Histological sections indicated minor developmental delay in the homozygotes, notably in the mid and hind brains (**Figure 1B**). Consistent with a complete loss of signalling activity (Pridans et al., 2013) and as reported previously in the context of analysis of a *Csf1r* hypomorphic mutation (Munro et al., 2020), the *Csf1r*^E631K/E631K^ embryos lacked detectable IBA1^+^ macrophages throughout the embryo apart from a small number of IBA1^+^ cells in the liver (**Figure 1C-F**). Compared to WT, there was an apparent 30-40% reduction (p<0.05) in IBA1^+^ cell populations in liver and throughout the *Csf1r*^E631K/+^ embryo. CSF1R^+^ macrophages are engaged in active clearance of apoptotic cells during embryonic development, notably between the digits, in the pharyngeal arches and in removal of expelled erythrocyte nuclei in the liver (Lichanska et al., 1999). However, despite the severe depletion of macrophages in the *Csf1r*^E631K/E631K^ mice, we saw no evidence of accumulation of pyknotic nuclei.

**Figure 1:**
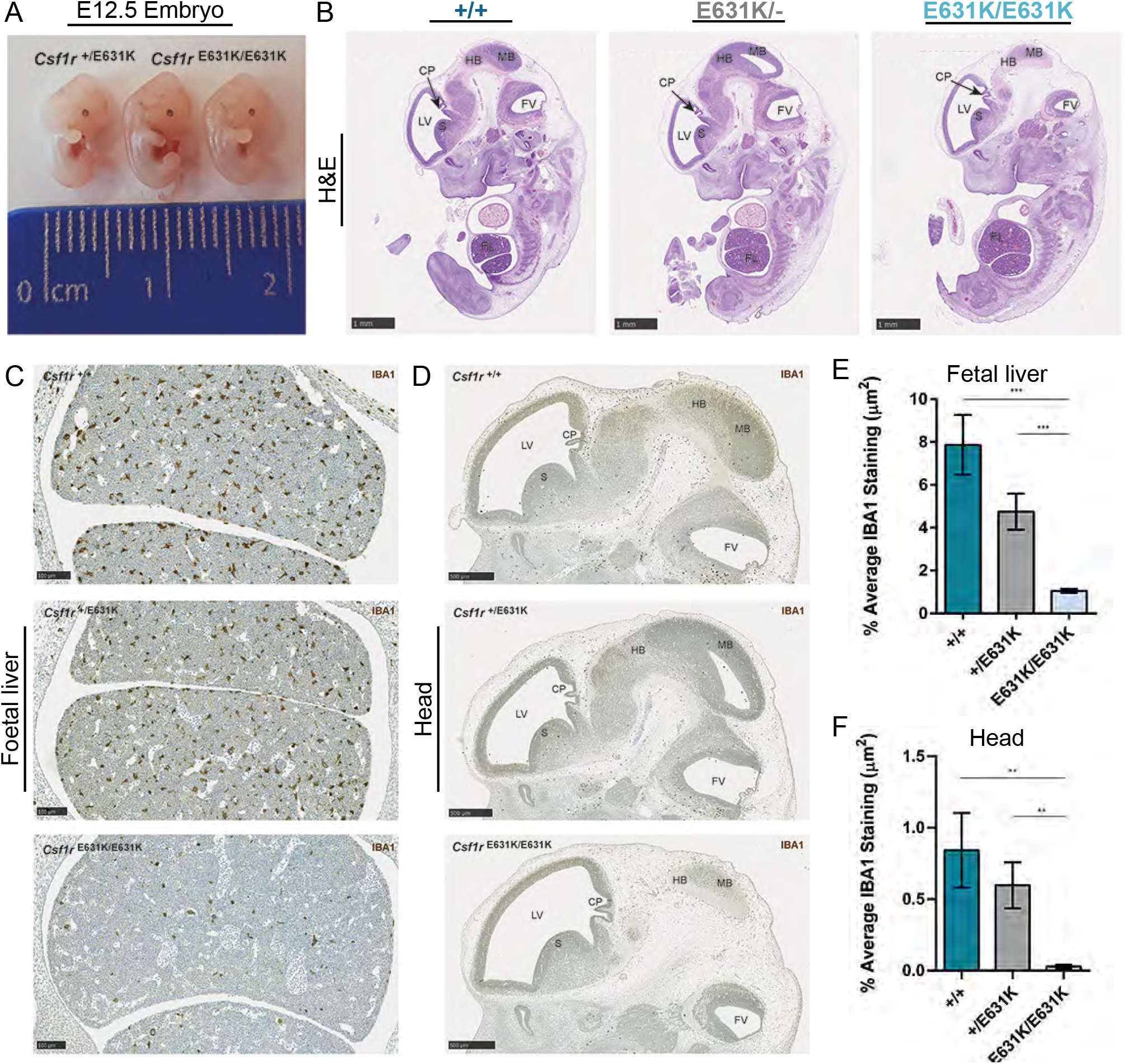
The effect of *Csf1r* E631K mutation on the mouse embryo. *Csf1r^E631K/+^* male and female mice were mated and pregnancies were terminated at 12.5dpc. Embryos were genotyped and processed for histology and detection of IBA1^+^ macrophages by immunohistochemistry as described in Materials and Methods. **(A)** Comparison of gross morphology of a *Csf1r^E631K/+^* embryo and two *Csf1r^E631K/E631K^* littermates. **(B)** Representative midline sections of each genotype stained with hematoxylin and eosin (H&E). Representative IBA1 staining in the **(C)** foetal liver and **(D)** E12.5 head of each genotype; LV = left ventricle, CP = choroid plexus, HB = hindbrain, MB = midbrain, FV = fourth ventricle, S = striatum, FL = foetal liver. Morphometric analysis of IBA1 immunolabelling in **(E)** foetal liver and **(F)** E12.5 head. Graphs shows mean +/-SEM. +/+ (n=3), +/E631K (n=4), E631K/E631K (n=6). ***p< 0.001; **p<0.01 via unpaired Student’s t-test.

### The impact of heterozygous Csf1r^E631K^ mutation on postnatal development of tissue macrophage populations

On the C57Bl/6JCrl background in Edinburgh, in heterozygous mating there were no live *Csf1r*^E631K/E631K^ pups. The number of *Csf1r*^E631K/+^ at weaning was not significantly different from expected 2:1 ratio of heterozygous:wild type. On the C57Bl/6JArc background in Australia, occasional *Csf1r*^E631K/E631K^ pups were born. The few live *Csf1r*^E631K/E631K^ mice weighed less than half their littermates by 3 weeks and had severely impaired skeletal development (**Figure 2A**). In the *Csf1r*^E631K/+^ x *Csf1r*^+/+^ matings, there was no significant loss of *Csf1r*^E631K/+^ pups prior to weaning. Like the *Csf1r* knockout on the C57Bl/6J genetic background (Erblich et al., 2011), the *Csf1r*^E631K/E631K^ pups lacked microglia, there was severe ventricular enlargement and the corpus callosum was almost undetectable (not shown). By contrast, the heterozygous mutant (*Csf1r*^E631K/+^) male mice had a small transient lag in postnatal growth (**Figure 2B**) but otherwise like the *Csf1r*^ΔFIRE/ΔFIRE^ hypomorphic line (Rojo et al., 2019) they developed normally, were healthy and fertile and showed no evident behavioural phenotype.

**Figure 2:**
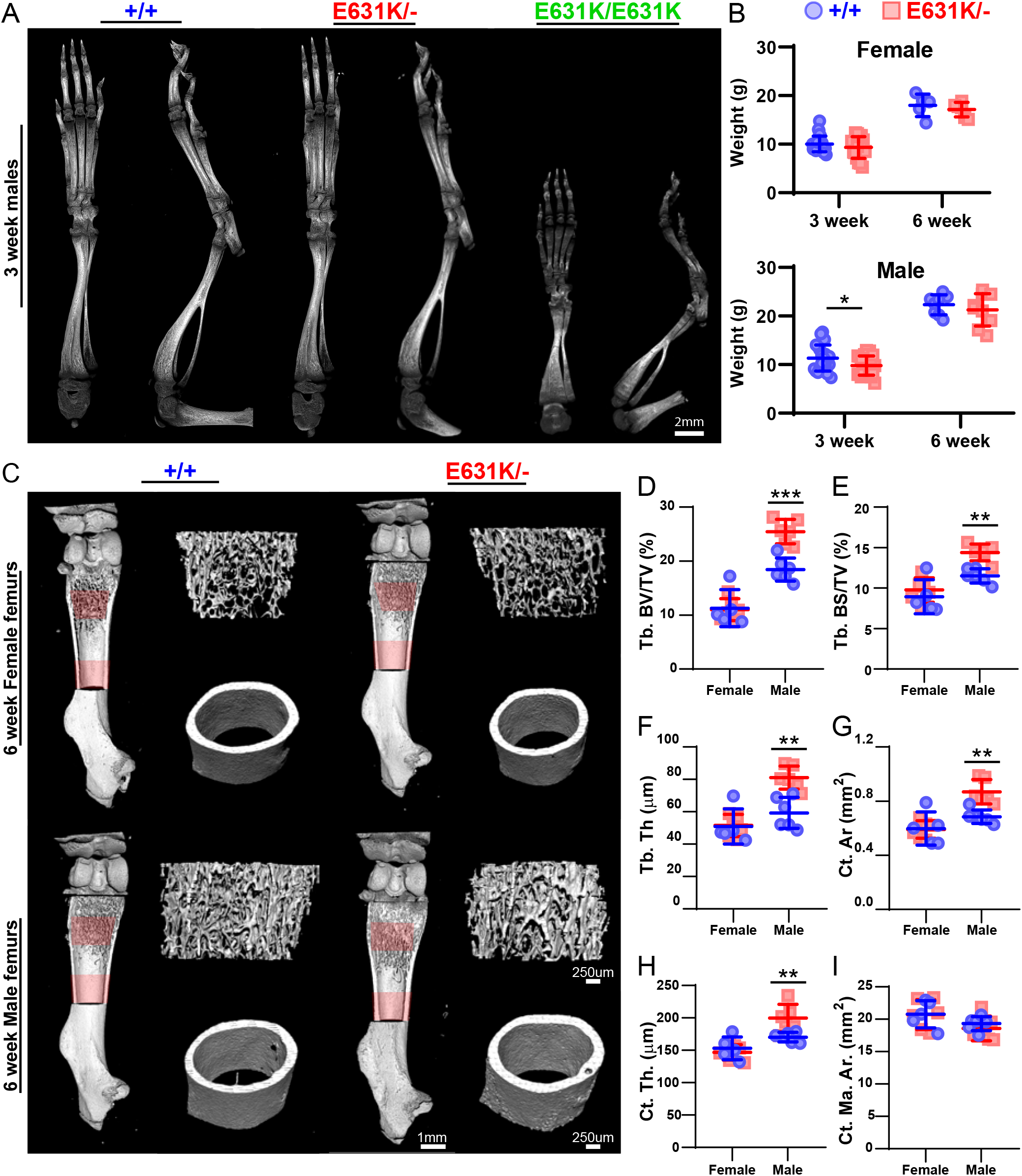
The effect of heterozygous *Csf1r* E631K mutation on bone development and postnatal growth. **(A)** Representative 3D reconstruction of the left hindlimb from micro-CT images of 3-week-old male *Csf1r^+/+^*, *Csf1r^E631K/+^, Csf1r^E631K/E631K^* mice. **(B)** Female and male mice weights were monitored and recorded at 3 and 6 weeks of age. **(C)** Representative 3D reconstruction of the femurs from micro-CT images of 6-week-old female and male *Csf1r^+/+^* and *Csf1r^E631K/+^* mice. Scans were performed in 10um slices and a depth of 1500um (or 150 slices) was analysed for both trabecular region (start identified as point of fusing of growth plates) and cortical region (start identified as last point of trabecular bone). Regions analysed are highlighted in red. **(D-I)** Micro-CT analysis of **(D)** percentage trabecular (Tb.) bone volume over tissue volume (BV/TV), **(E)** Tb Bone surface to volume ratio (BS/TV), **(F)** Tb thickness (Tb. Th), **(G)** cortical (Ct.) bone area (Ar), **(H)** Ct. thickness (Ct. Th.), and **(I)** marrow area in the cortical region analysed (Ct. Ma. Ar.). Data are derived from 4-6 mice of each sex for each genotype at 6 weeks of age. Individual data points with mean and standard deviation are presented in each graph. Statistical analysis was performed using unpaired Student’s t-test: * = P<0.05, ** = P<0.001, *** = P<0.0001.

The definitive phenotype of *Csf1* and *Csf1r* mutations in mice and rats is osteoclast deficiency and osteopetrosis (Chitu and Stanley, 2017; Hume et al., 2020). μCT analysis of adult male and female mice confirmed the substantially lower trabecular bone density and trabecular volume (**Figure 2C-D**) that was described previously in female C57Bl/6J mice (Sauter et al., 2014). The *Csf1r*^E631K/+^ male mice showed a significant increase in trabecular bone density and cortical bone thickness compared to controls whereas the mutation did not overcome the osteoporosis seen in females (**Figure 2C-I**).

Macrophage populations of the mouse and rat expand substantially in the postnatal period. Organs grow rapidly and macrophage density in each organ also increases as evident from the increase in relative expression of macrophage-expressed transcripts including *Csf1r* (Summers and Hume, 2017). The postnatal expansion of the resident mononuclear phagocyte populations is associated with a postnatal increase in *Csf1* mRNA in most organs and is CSF1R-dependent as shown by the impacts of both the *Csf1*^op/op^ and *Csf1r* mutations and the effect of postnatal treatment with anti-CSF1 antibody (Cecchini et al., 1994; Keshvari et al., 2021; Rojo et al., 2019; Summers and Hume, 2017; Wei et al., 2005). We predicted that a dominant-negative effect of the *Csf1r*^E631K^ allele would compromise and delay this postnatal resident tissue macrophage expansion. As an initial screen for the impact of the mutation on CSF1R-dependent macrophage proliferation we examined wholemounts using the C*sf1r-*EGFP reporter transgene at 3 weeks and 7 weeks of age. At 3 weeks in the *Csf1r*^E631K/+^ mice, *Csf1r-*EGFP positive macrophages were either absent or greatly reduced in every organ examined. **Figure 3** shows diverse examples: the skin (both ear and foot), adrenal gland, lung, liver, adipose, pancreas and testis. By 7 weeks of age (**Figure S2**) most of these tissue macrophage populations appeared indistinguishable between *Csf1r*^E631K/+^ and *Csf1r*^+/+^ littermates. On close examination, in areas such as pancreas and adipose (where the stellate morphology of mature macrophages is more obvious), many of the *Csf1r*-EGFP^+^ cells appeared round and monocyte-like suggesting population of the niche was ongoing in the mutant mice. Notably, despite the clear reduction in macrophage density at 3 weeks, the regular spacing that is evident in all tissue-resident populations (Hume et al., 2019) was established. One surprising feature was an apparent difference between Langerhans cell (LC) populations in the ear and footpad. The LC population of the footpad remained absent at 7 weeks, whereas the normal LC density was evident in the ear. To corroborate and quantify the loss of macrophages evident in whole mount imaging, we located IBA1^+^ macrophages in the cortex of the kidney, where *Csf1*^op/op^, *Csf1r*^-/-^ (Dai et al., 2002) and *Csf1r*^ΔFIRE/ΔFIRE^ mice (Rojo et al., 2019) are almost completely macrophage-deficient (**Figure 4**A). At 3 weeks, *Csf1r*^E631K/+^ mice had only 30% of the number of renal cortical interstitial IBA1^+^ cells detected in their WT littermates, 53 % by 7 weeks, and by 9 weeks this was increased to 72% (**Figure 4B**). Despite the delay in macrophage population there was no apparent impact on glomerular or tubular development (**Figure 4A**).

**Figure 3.**
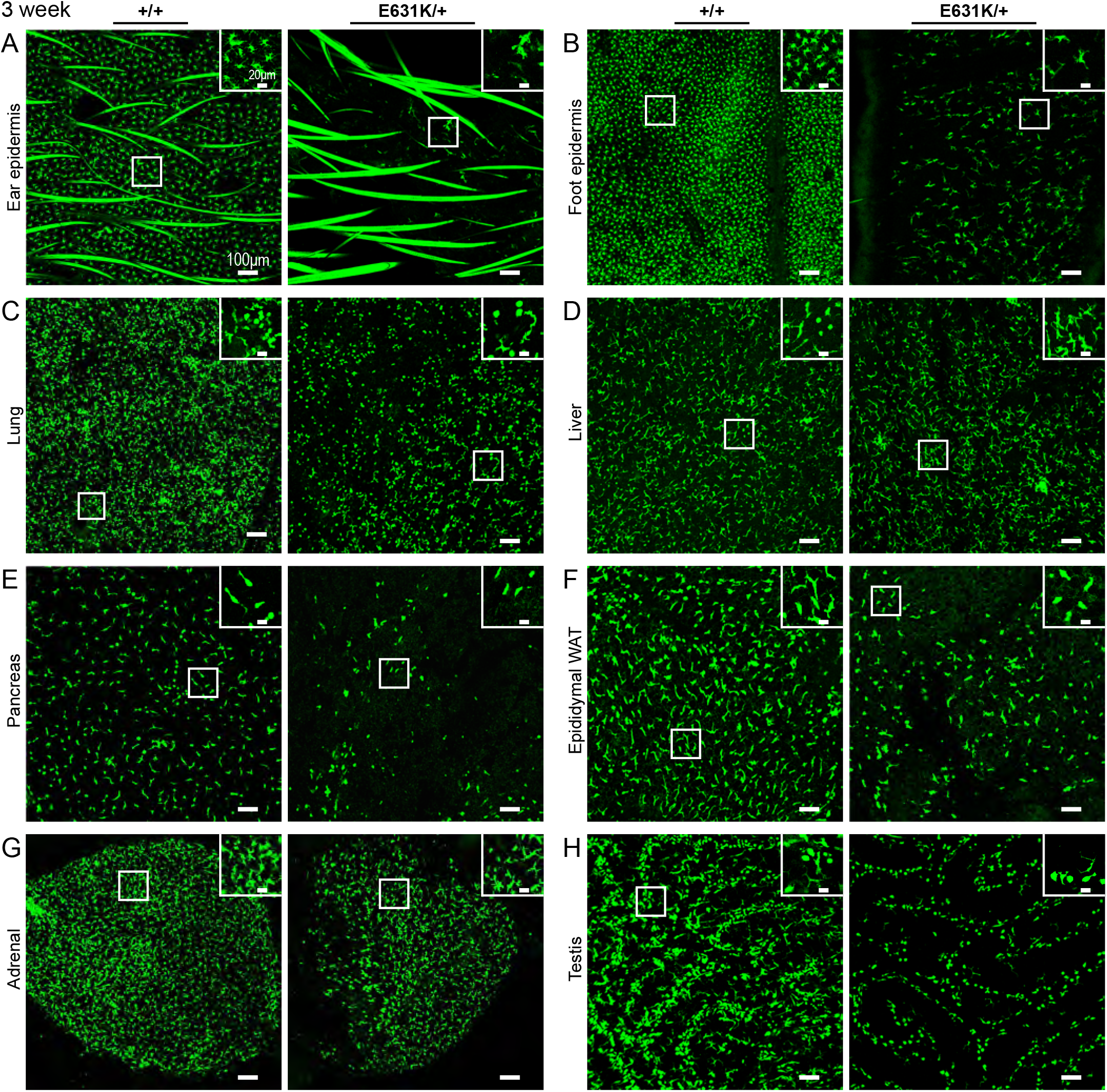
The effect of heterozygous *Csf1r* E631K mutation on resident tissue macrophage populations at 3 weeks of age. Tissues were harvested from 3-week-old male *Csf1r^+/+^* and *Csf1r^E631K/+^* littermates, each also *Csf1r-*EGFP transgenic. Whole-mounted tissues were placed in PBS on ice and imaged directly within 2–3 h using an Olympus FV3000 microscope. Images show the same depth of maximum intensity projections of the tissues indicated in **(A)** to **(H)**. The lung **(C)** and liver **(D)** projections include stellate subcapsular populations (see insets). Note comparable images from 7-week-old males of both genotypes are shown in **Fig. S2**. Images are representative of at least 3 mice from each genotype.

**Figure 4.**
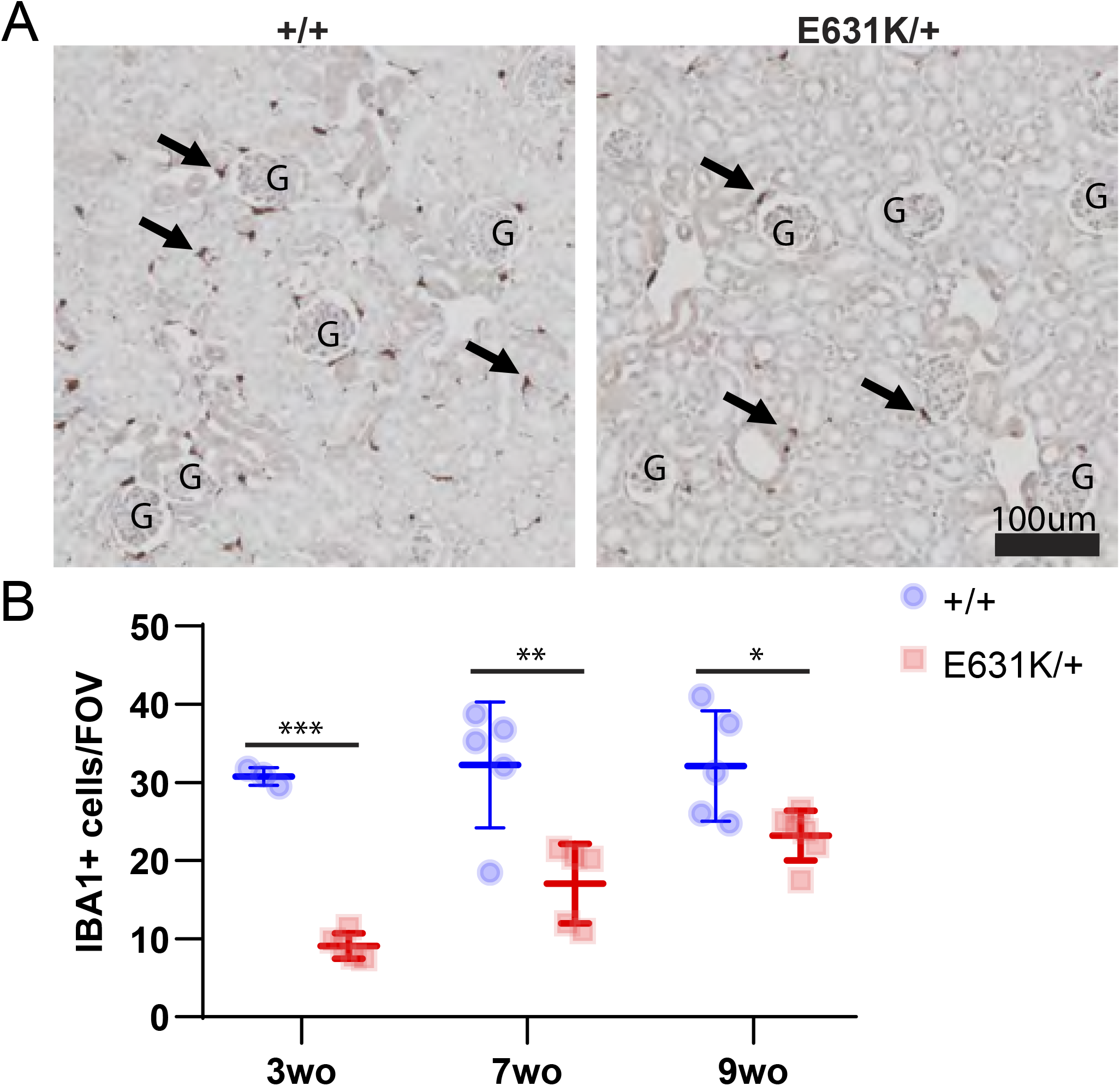
Age-dependent effect of heterozygous *Csf1r* E631K mutation on resident renal macrophage populations. Kidneys from *Csf1r^+/+^* and *Csf1r^E631K/+^* littermates were isolated at each of the ages indicated, fixed and embedded. **(A)** Representative IBA1 staining from each genotype at 3 weeks of age; G = glomerulus, arrows = IBA1+ cells. **(B)** The number of IBA1+ cells in the cortex were quantified by averaging the number of cells from 4 FOV’s (area = 0.25μm^2^) per animal. Data are derived from 4-6 mice for each genotype at 3, 7, and 9 weeks of age. Individual data points with mean and standard deviation are presented in each graph. Statistical analysis was performed using unpaired Student’s t-test: * = P<0.05, ** = P<0.001, **** = P<0.0001.

**Figure 5A** shows flow cytometry analysis of bone marrow (BM) progenitor and monocyte populations in 3-week-old animals. In *Csf1r*^ΔFIRE/ΔFIRE^ mice, CSF1R was undetectable in BM or blood based on binding of anti-CD115 or CSF1-Fc, yet there was no detectable change in BM or blood monocyte profiles (Rojo et al., 2019). Monocyte numbers are also unaffected by anti-CSF1R treatment in adults (MacDonald et al., 2010). Hence, although CSF1 treatment can drive monocyte proliferation *in vitro* and *in vivo*, it is not absolutely required for monocyte production. We analysed hematopoietic stem and progenitor, committed progenitor and monocyte populations using surface markers as previously described (Grabert et al., 2020). **Figure 5A/B** shows representative FACS profiles and gating. There was no significant effect of *Csf1r*^E631K/+^ mutation on the abundance of stem and progenitor cells, committed precursors or mature myeloid cell populations compared to controls. However, the level of expression of CSF1R on individual cells detected using anti-CD115 was reduced in *Csf1r*^E631K/+^ BM populations with the greatest impact in progenitors (**Figure 5C-F**). **Figure 5G-I** shows analysis of peritoneal populations. Whereas the relative abundance of F4/80^hi^ peritoneal macrophages was unchanged in *Csf1r*^E631K/+^ mice, there was selective loss of the F4/80^lo^ subset (**Figure 5L**). These cells are monocyte-derived and may serve as precursors for the slow replacement of embryo-derived macrophages (Bain et al., 2016). In both peritoneal macrophage populations surface CD115 was detectable and clearly reduced in both populations in the *Csf1r*^E631K/+^ mice (**Figure 5J**). However, precise quantitation was compromised by autofluorescence.

**Figure 5.**
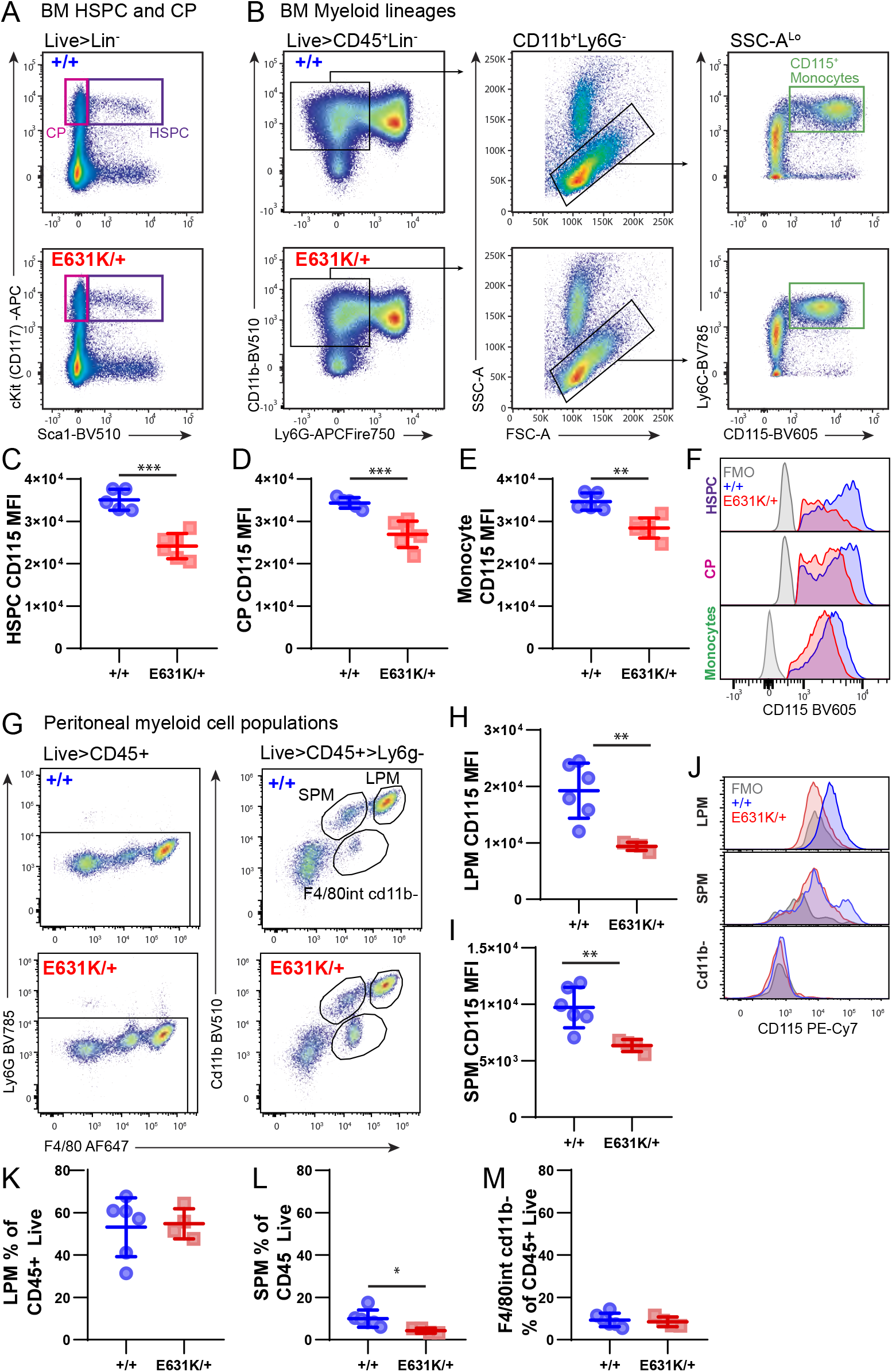
The heterozygous *Csf1r* E631K mutation on bone marrow and peritoneal macrophage populations and CSF1R expression. Bone marrow and peritoneal cells were isolated from *Csf1r^+/+^* and *Csf1r^E631K/+^* littermates at 3 weeks of age. **(A) – (F)** Bone marrow (BM) cKit^+^Sca1^+^ hematopoietic stem and progenitor cells (HSPC), cKit^+^Sca1^-^committed progenitors (CP), and mature myeloid cells were analysed by Flow Cytometry and gated as previously reported (Grabert et al., 2020). Representative flow cytometry plots of **(A)** HSPC (purple gate) and CP (red gate), and **(B)** of mature myeloid cells of WT (+/+) and E631K heterozygous (E631K/-) mice. Note the left shift of anti-CD115 staining in monocytes in the *Csf1r^E631K/+^* mice (upper right Panel B). CD115 mean fluorescence intensity (MFI) for **(C)** HSPC, **(D)** CP and **(E)** Ly6C+CD115+ monocytes. **(F)** Overlaid representative histograms (normalized to mode) for each genotype and the fluorescence minus 1 (FMO) control. Note that the detection of CD115 in progenitors is heterogeneous and the effect of the *Csf1r^E631K/+^* mutation may be greater than evident from the MFI. **(G)** – **(K)** Flow cytometry analysis of CD45^+^ peritoneal (PT) cells from each genotype. **(G)** Representative flow cytometry plots of PT cells stained for F4/80 and CD11b. CD115 MFI in **(H)** large peritoneal macrophages (LPM) and **(I)** small peritoneal macrophages (SPM). **(J)** Overlaid representative histograms (normalized to mode) for CD115 staining for each population and each genotype and the FMO control. Note that the FMO control in this case is high because of significant autofluorescence and CD115 staining in the SPM is heterogeneous. Proportional quantification of **(K)** LPM, **(L)** SPM and **(M)** F4/80^int^CD11b^-^ cells (each of total CD45^+^ cells). Data are derived from 4-6 mice for each genotype at 6 weeks of age. Individual data points with mean and standard deviation are presented in each graph. Statistical analysis was performed using unpaired Student’s t-test: * = P<0.05, ** = P<0.001, *** = P<0.0001.

### The impact of heterozygous Csf1r^E631K^ mutation on brain microglial populations, motor functions and pathology

Imaging using the *Csf1r*-EGFP reporter indicated that unlike *Csf1r*^ΔFIRE/ΔFIRE^ mice, the *Csf1r*^E631K/+^ mice were not globally deficient in microglia, but there was a clear phenotype. **Figure 6A** shows a comparison of *Csf1r*-EGFP in cortical microglia in WT and *Csf1r*^E631K/+^ littermates at 7 weeks of age. Aside from the decrease in the number of microglial cell bodies in any field, there was an obvious reduction of dendritic arborisation of the membrane processes in the mutant. To document these changes more thoroughly, we performed IBA1 staining on multiple brain regions at different ages (**Figure 6B-G**). These analyses were carried out independently in both Brisbane and Edinburgh where the mutations were on subtly different C57BL/6J backgrounds. The *Csf1r*^E631K/+^ had 20-40% reduced IBA1^+^ microglial numbers in all brain regions measured in juveniles (3 weeks), young adults (7-9 weeks) and aged (43 weeks) mice (**Figure 6D**). Note a downward trend in microglial number with age independent of *Csf1r* genotype. The 9 weeks data generated independently in Edinburgh for cerebellum and forebrain is shown in **Fig. S3** and **S4.** CSF1-dependent microglia have been attributed functions in development and turnover of Purkinje cells (Kana et al., 2019; Marin-Teva et al., 2004) but there was no evidence of any phenotype in the cerebellum, despite reduced numbers and apparent ramification of microglia in both white and grey matter.

**Figure 6.**
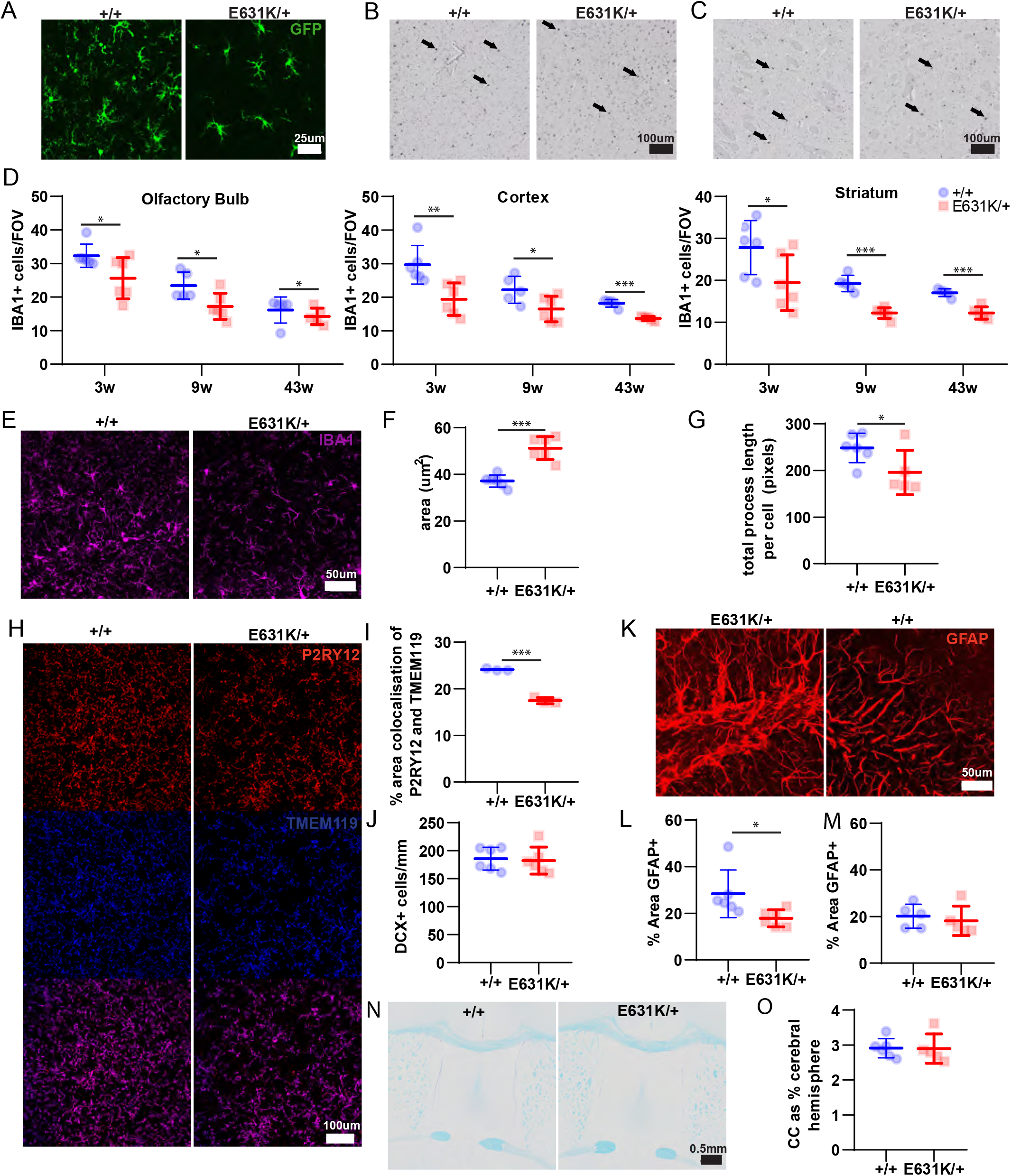
The effect of heterozygous *Csf1r* E631K mutation on microglia and other cell populations in the brain. Brain tissues were isolated from cohorts of *Csf1r^+/+^* and *Csf1r^E631K/+^* mice at 3-, 7-, 9 and 43 weeks of age, fixed and processed for immunohistochemistry as detailed in Materials and Methods. **(A)** Representative MIPs of confocal z-stack series of wholemount cortex from 7-week-old mice that were transgenic for Csf1r-EGFP. Images show the same depth of maximum intensity projections. **(B)** Representative IBA1 staining in the cortex, and striatum **(C)** of 9-week-old mice. **(D)** Quantitative analysis of IBA1+ cells in the olfactory bulb, cortex, and striatum of 3-, 9-, and 43-week-old mice. The brains of 7-week-old mice were examined with IFC **(E, H, K)**. **(E)** MIP of confocal z-stack series showing IBA1+ cells in the dentate gyrus. **(F)** Quantitative analysis was performed on microglial cell body area (average of 20 cells/animal), and **(G)** the average total process length per cell (average of 3 FOVs per animal). **(H)** Representative MIPs of P2RY12 and TMEM119, and colocalised staining in the cortex. **(I)** Quantitative analysis of the percentage area of colocalised P2RY12 and TMEM119. **(J)** Quantitative analysis of the number of DCX+ cells/mm of DG (average of 3 DGs/animal). **(K)** Representative MIP of confocal z-stack series showing GFAP staining in the dentate gyrus. **(L)** Quantitative analysis was performed on the % area GFAP+ staining (average of 3 areas) of 7-week-old brains using IFC, and **(M)** 43-week-old brains using IHC. **(N)** Representative luxol fast blue with cresyl fast violent counter stain in 9-week-old brains. **(O)** Quantitative analysis of the area of half of the corpus callosum as a percentage area of the cerebral hemisphere at 9 weeks. n=5-6/group. Graphs show mean and standard deviation. Statistical analysis was performed using unpaired Student’s t-test: *, **, ***, **** = p <0.005, 0.01, 0.001, 0.0001.

To quantify the effect on microglial arborisation, we analysed images of individual microglia from 7-week-old brains using ImageJ. **Figures 6 F/G** confirm that microglia in the *Csf1r*^E631K/+^ mice had an increased cell body area, and greatly reduced ramification of membrane processes. Altered morphology is considered an indication of reactive microglia, and is a feature of human ALSP (Kempthorne et al., 2020; Tada et al., 2016). In a model of Cre-induced heterozygous mutation of *Csf1r* (Arreola et al., 2021) microglial dyshomeostasis was evident from reduced expression of microglia-enriched markers such as P2RY12, but this model is also heterozygous for a *Cx3cr1* mutation. **Figure 6H** shows immunolocalisation of homeostatic markers P2RY12 and TMEM119 in *Csf1r*^+/+^ and *Csf1r*^E631K/+^ cortex. Localisation of these markers confirmed the reduced density (**Figure 6I**) of microglia seen with IBA1 but the staining intensity and punctate distribution of these markers on individual cells was unaffected by the *Csf1r* mutation.

Microglia have been attributed many functions in postnatal brain development through interactions with neurogenic progenitors, neurons, astrocytes and oligodendrocytes (Han et al., 2021; Matejuk and Ransohoff, 2020; Prinz et al., 2019). Microglia interact with and regulate the function of astrocytes and astrogliosis is a common feature of neuroinflammatory diseases including ALSP (Han et al., 2021; Matejuk and Ransohoff, 2020). We analysed the distribution of astrocytes and doublecortin (DCX)-positive neurogenic progenitors in the hippocampus of 7-week-old *Csf1r*^E631K/+^ and control mice (**Figure 6J-M**). Within the dentate gyrus, the apparent density of GFAP-staining was almost 40% reduced in 7-week-old *Csf1r*^E631K/+^ mice whereas there was no discernible difference in the number of DCX^+^ neurons (**Figure 6J-M**). In the 43-week-old *Csf1r*^E631K/+^ mice, there was no longer any difference in GFAP density compared to controls.

The reduced microglial density and altered morphology observed in the *Csf1r*^E631K/+^ mice is consistent with the human disease (Kempthorne et al., 2020; Tada et al., 2016) but the opposite of the increased microglial density reported in *Csf1r*^+/-^ mice on the C57BL/6J background (Arreola et al., 2021; Chitu et al., 2020; Chitu et al., 2015). The *Csf1r*^ΔFIRE/+^ mice also have a 50% reduction in *Csf1r* mRNA in the juvenile brain, but comparable microglial density as indicated by expression of a suite of microglia-associated transcripts (Rojo et al., 2019). However, as shown in **Figure S5,** we were able to confirm that the 50% loss of microglial CSF1R was associated with a 2-fold increase in microglial density in hippocampus, motor cortex and corpus callosum in 6-month-old C57Bl/6J *Csf1r*^ΔFIRE/+^ mice compared to WT. There was also a marginal increase in GFAP+ astrocytes in the hippocampus.

Many ALSP patients present initially with sensorimotor deficiencies (Chitu et al., 2021; Konno et al., 2018; Konno et al., 2017) and CSF1R signalling has been directly implicated in neuropathic pain (Saleh et al., 2018). To seek evidence of these symptoms in the *Csf1r*^E631K/+^ mice, we performed a wide range of sensorimotor tests. As shown in **Figure 7,** we could not demonstrate any significant impacts of the *Csf1r*^E631K/+^ genotype on gross motor function or mechanical paw withdrawal thresholds measured at 10 months of age.

**Figure 7.**
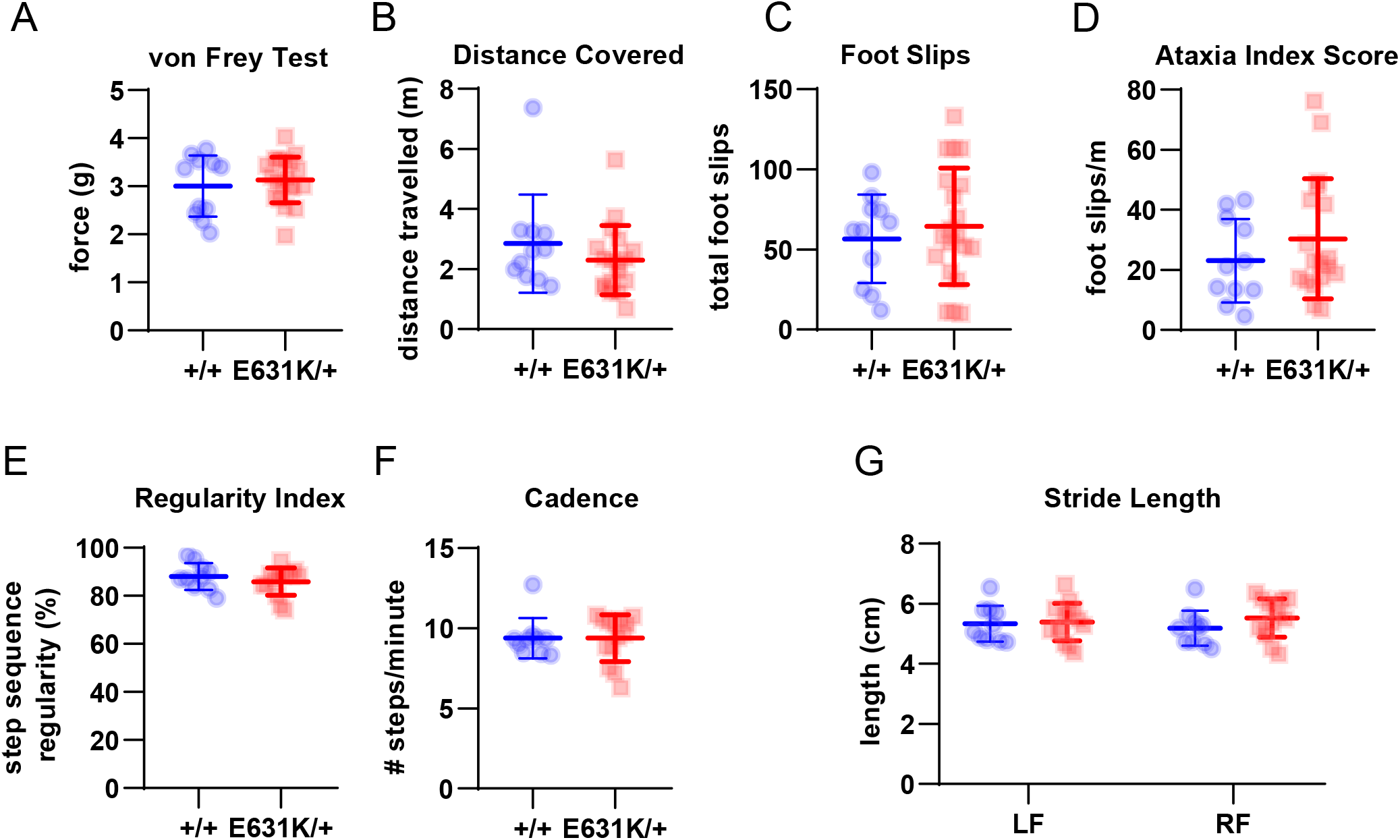
The effect of *Csf1r* genotype on sensorimotor parameters in aged mice. A large cohort of 43-week-old *Csf1r^+/+^* and *Csf1r^E631K/+^* male and female mice was subjected to a range of sensorimotor tests as described in Materials and Methods. **(A)** Quantification of the Von Frey test as a measurement of mechanical allodynia in rodents. The vertical axis shows the threshold force at which the mice withdraw the paw, a measure of pain sensitivity. **(B)** – **(D)** Quantification of the parallel rod floor test as a measure of balance and coordination, quantified as **(B)** distance covered, **(C)** number of foot slips, and **(D)** ataxia index score. **(E)** – **(G)** Quantification of the CatWalk XT gait analysis showing results for **(E)** step regularity index, **(F)** cadence, or **(G)** stride length of either the left front (LF) or right front (RF) paw. Data are derived from 10-14 mice for each genotype at 43 weeks of age. Individual data points with mean and standard deviation are presented in each graph. Statistical analysis was performed using unpaired student’s t-test with no statistical significance between genotypes.

To further test for age-dependent effects of *Csf1r*^E631K/+^ genotype we also examined an even older cohort (15 months). No periventricular calcification or evidence of pathology was detected by magnetic resonance imaging (**Figure S6A**) or upon histological examination of multiple brain regions (not shown). Chitu *et al*. (Chitu et al., 2020; Chitu et al., 2015) emphasised the development of microgliosis in the white matter of *Csf1r*^+/-^ mice as a model of ALSP. IBA1 staining of these aged brains revealed occasional clusters of reactive microglia and apparent microglial heterogeneity in striatum and corpus callosum (not shown) but with equal prevalence in *Csf1r*^+/+^ and *Csf1r*^E631K/+^ mice. The absolute differences in overall IBA1^+^ cell density seen in younger animals were no longer evident (**Figure S6B**). Confirming this conclusion, disaggregation of the aged brains yielded similar numbers of microglia with unchanged profiles of fluorescence intensity of microglial markers CD11B, CD45 and P2RY12 (**Figure S7**).

In summary, the data indicate that heterozygous *Csf1r* kinase-dead mutation in mice has a dominant inhibitory effect on development of CSF1R-dependent microglia and selected tissue macrophage populations.

### Dominant repression of CSF1 responsiveness in the periphery in Csf1r^E631K/+^ mice

Whereas there was a relatively mild phenotype in the brain, the heterozygous kinase-dead mutation clearly impacted development of CSF1R-dependent resident macrophage populations in the periphery (**Figure 3**). We therefore examined whether these impacts could be linked directly to a loss of CSF1-responsiveness. BM cells respond to addition of CSF1 in liquid culture to generate confluent cultures of adherent macrophages within 5-7 days. In *Csf1r*^ΔFIRE/ΔFIRE^ mice which lack CSF1R expression in bone marrow progenitors, this response was abolished (Rojo et al., 2019). Consistent with the loss of CSF1R in progenitors (**Figure 5**), BM cells isolated from *Csf1r*^E631K/+^ mice were almost completely deficient in proliferation and differentiation response to CSF1 *in vitro* whereas the response to CSF2 (GM-CSF) was unaffected (**Figure 8A**).

**Figure 8.**
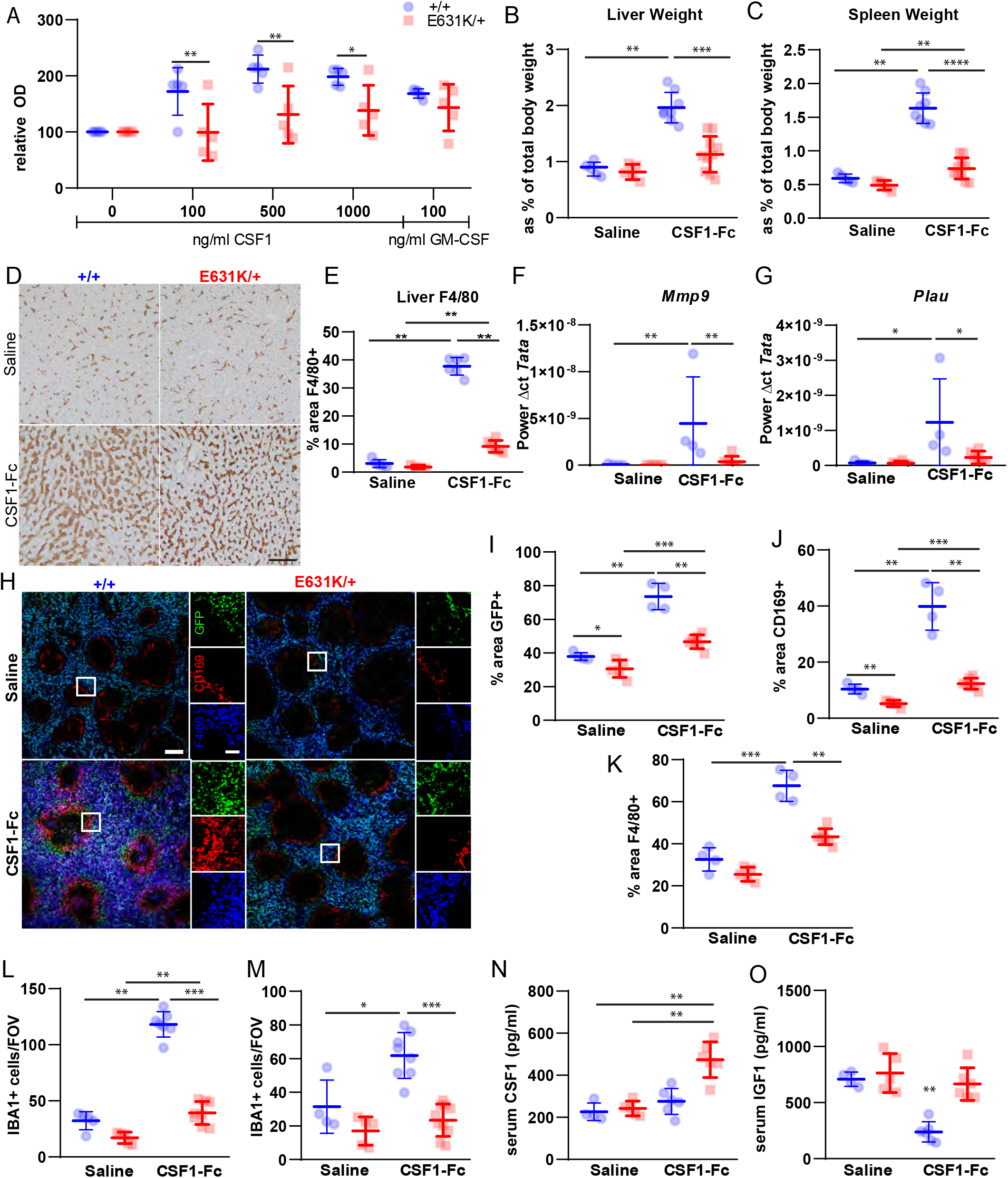
The effect of heterozygous *Csf1r* E631K mutation on responses to CSF1 *in vitro* and *in vivo*. *In vitro response*. **(A)** Bone marrow cells from *Csf1r^+/+^* and *Csf1r^E631K/+^* mice were cultured for 7 days in recombinant human CSF1 at the concentrations indicated or in mouse GM-CSF (100ng/ml). Live cells were quantified using the Resazurin assay as described in Materials and Methods and results are expressed as optical density relative to the unstimulated cultures in each case. n= 5/group; 4 technical replicates per animal. Two-way ANOVA with multiple comparisons was performed: *, ** = p <0.05, 0.01 *in vivo response*. Mice were injected with CSF1-Fc (5mg/kg) or saline control on each of 4 successive days and euthanised on day 5. **(B)** The liver and **(C)** spleen of all animals were weighed following acute in vivo CSF1. **(D)** Representative images of F4/80 IHC staining of liver sections from all treatment groups. Scale bar = 50μm. **(E)** The % area stained for F4/80 in liver sections was quantified (average of 4 areas/animal). qPCR analysis shows upregulation of CSF1R target genes including **(F)** *Mmp9* and **(G)** *Plau*. **(H)** Representative images of GFP, CD169, and F4/80 staining of spleen sections from all treatment groups. Scale bar in main image = 200μm, in insert = 50μm. The % area stained for **(I)** GFP, **(J)** CD169, and **(K)** F4/80 in spleen sections was quantified (average of 2 depths/animal, with least 50% of each spleen on the sagittal sections analysed). Quantitative analysis of IBA1+ cells in the **(L)** kidney and **(M)** heart. **(N)** Serum CSF1 and **(O)** IGF1 were measured. n= 4-9 per group. Mann-Whitney: *, **, ***, **** = p <0.05, 0.01, 0.001, 0.0001.

To determine whether the CSF1 resistance *in vitro* was also manifest *in vivo*, we examined the response to administration of exogenous CSF1. Treatment of mice with a pig CSF1-Fc fusion protein caused a monocytosis, expansion of tissue macrophage populations through monocyte recruitment and local proliferation and consequent hepatosplenomegaly (Gow et al., 2014). Here we used a human CSF1/mouse Fc protein that has similar biological activity to the pig protein and was prepared for pre-clinical evaluation. **Figure 8 B-O** compares the responses of *Csf1r*^E631K/+^ mice and *Csf1r*^+/+^ controls to acute administration of a maximal dose of CSF1-Fc on 4 successive days with analysis on day 5. The monocytosis, increased size of the liver and spleen, expansion of F4/80^+^ macrophage populations observed in controls in response to CSF1-Fc was almost entirely absent in *Csf1r*^E631K/+^ mice (**Figure 8B-E**). Analysis of liver mRNA confirmed the induction of CSF1R target genes, *Plau* and *Mmp9* (Gow et al., 2014), in controls that was undetectable in *Csf1r*^E631K/+^ mice (**Figure 8F, G**). One novel circulating biomarker of the response to CSF1-Fc is the somatic growth factor, insulinlike growth factor 1 (IGF1), which was reduced to almost undetectable levels in serum from treated *Csf1r*^+/+^ mice (**Figure 8O**), likely associated with proliferative expansion of the liver (Gow et al., 2014). IGF1 in serum was unaffected by CSF1-Fc in the *Csf1r*^E631K/+^ mutant (**Figure 8O**). Treatment with human CSF1-Fc effectively competes with the endogenous ligand and causes a transient increase in circulating mouse CSF1 that is resolved by day 5 due to expansion of tissue macrophages and consequent CSF1 clearance. Consistent with the lack of efficacy in *Csf1r*^E631K/+^ mice, endogenous CSF1 remained elevated in these mice on day 5 (**Figure 8N**).

The CD169^+^ marginal zone macrophage population in spleen is known to be especially CSF1/CSF1R-dependent in both mouse and rat (Pridans et al., 2018; Witmer-Pack et al., 1993). Consistent with the selective loss of CSF1R-dependent macrophages, this population remained greatly reduced in untreated adult *Csf1r*^E631K/+^ spleen whereas the F4/80^+^/*Csf1r*-EGFP^+^ cells of the red pulp were less affected (**Figure 8H-K**). Splenomegaly in CSF1-Fc treated *Csf1r*^+/+^ mice was associated with both expansion of the marginal zone CD169+ populations and extensive expansion of the CD169+ populations in the red pulp. Both these responses were prevented in the *Csf1r*^E631K/+^ mice (**Figure 8J**). As noted above, *Csf1r*^E631K/+^ mice had reduced numbers of macrophages in kidney and heart even as adults. CSF1-Fc treatment expanded the IBA1^+^ macrophage populations in both organs in *Csf1r*^+/+^ mice but had no effect in the heterozygous *Csf1r*^E631K/+^ mice (**Figure 8L, M**).

## Discussion

We have analysed the effect of a germ line kinase-dead *Csf1r* mutation (*Csf1r*^E631K^) associated with human ALSP on the mouse mononuclear phagocyte system. Homozygous E631K mutation phenocopied the impact of a homozygous null mutation (*Csf1r*^-/-^ (Chitu and Stanley, 2017; Erblich et al., 2011)) confirming that the mutation abolishes signalling activity. Like the knockout mutation on the C57BL/6J background, homozygous mutant (*Csf1r*^E631K/E631K^) embryos lacked macrophages (**Figure 1**). The few pups that survived embryonic development and were born had severe postnatal growth retardation and hydrocephalus.

The postnatal developmental phenotype in *Csf1r*^E631K/+^ mice is consistent with a partial loss of CSF1R activity. *Csf1r*^E631K/+^ mice had a reduced postnatal growth rate, mild male osteopetrosis and delayed development of tissue macrophage populations (**Figure 2, 3, S3**). The tissues impacted by *Csf1r*^E631K/+^ mutation include the lung, where CSF2 (GM-CSF) is the major growth factor required for development of alveolar macrophages and homeostasis (Guilliams et al., 2013) but CSF1 is required for postnatal expansion of resident macrophages ((Jones et al., 2014) and references therein). The CD169^+^ marginal zone macrophages of spleen, which are entirely depleted in both *Csf1*^op/op^ mice and *Csf1r*^-/-^ rats (Pridans et al., 2018; Witmer-Pack et al., 1993) remained reduced even in adult *Csf1r*^E631K/+^ mice. The regular distribution of *Csf1r-*EGFP^+^ cells in every tissue in juvenile *Csf1r*^E631K/+^ mice despite the reduced density (**Figure 3**) supports a model in which macrophages occupy territories that are established by mutual repulsion rather than precisely-defined spatial niches (Hume et al., 2019).

The analysis of the response to CSF1 in BM cells *in vitro* provides unequivocal evidence that the disease-associated mutation does indeed have a dominant-negative effect on CSF1R signalling when expressed at normal physiological levels in the natural target cells. The level of surface CSF1R detected with anti-CD115 antibody was reduced in bone marrow progenitors in *Csf1r*^E631K/+^ compared to *Csf1r*^+/+^mice. Whereas disease-associated mutations did not prevent expression of CSF1R on the cell surface in transfected factor-dependent BaF3 cells (Pridans et al., 2013), the most common human mutation (I794T) compromised ectodomain shedding of CSF1R over-expressed in HEK293 cells (Wei et al., 2021). A dominant impact of the mutant protein on stability of the wild-type CSF1R cannot be excluded. However, the selective loss of CSF1R in marrow progenitors compared to monocytes (**Figure 5F**) might also be a consequence of a signalling defect, since *Csf1r* mRNA and protein expression is itself CSF1-inducible in progenitors and increases during differentiation (Grabert et al., 2020; Tagoh et al., 2002).

Regardless of the underlying mechanism, the *Csf1r*^E631K/+^ mice are not entirely CSF1R-deficient, since all CSF1R-dependent resident macrophages that can be depleted rapidly by anti-CSF1R (MacDonald et al., 2010) are present in adults. However, adult *Csf1r*^E631K/+^ mice were CSF1 resistant *in vivo*. Consistent with previous studies with pig CSF1-Fc (Gow et al., 2014; Stutchfield et al., 2015), the human-mouse CSF1-Fc used here promoted monocytosis, proliferative expansion of the liver and spleen and increased macrophage populations in multiple organs. We show here that CSF1-Fc treatment is associated amongst other impacts with a rapid fall in circulating IGF-1. We extended the earlier data (Gow et al., 2014) to demonstrate expansion of macrophage populations in the heart and kidney (organs where tissue macrophages are particularly CSF1R-dependent (Rojo et al., 2019) and the novel observation that CSF1 selectively expands CD169^+^ populations in both the marginal zone and red pulp of spleen. Mechanistic studies of the pleiotropic effects of CSF1-Fc administration are ongoing. The key finding herein is that all of these responses to CSF1-Fc administration were reduced or abolished in the *Csf1r*^E631K/+^ mice.

*Csf1r*^E631K/+^ mice had 20-40% reduced numbers of microglia in embryos, juveniles and adults in all regions of the brain. The more striking impact was reduced spreading and ramification which may reflect the well-studied ability of CSF1 to promote membrane ruffling, filipodia formation and spreading on a substratum (Stanley and Chitu, 2014). These mouse microglial phenotypes closely resemble those reported in human ALSP patients (Kempthorne et al., 2020; Tada et al., 2016). We have also generated mice with the most common human ALSP-associated mutation, *Csf1r*^I794T^, which is also signalling-deficient when expressed in Ba/F3 cells (Pridans et al., 2013). Preliminary analysis of these mice supports the dominant-negative model and partial microglial deficiency described in *Csf1r*^E631K/+^ mice. The microglial difference between *Csf1r*^E631K/+^ and *Csf1r*^+/+^ mice was no longer evident at 15 months. However, the data in **Figure 6** indicate that this convergence is due to a progressive reduction in microglial density in the controls rather than recovery in the mutant. There was no evidence of functional deficits or overt brain pathology resembling the human disease. The delay in microglial population and phenotypic modulation in younger animals was associated with a delay in astrocyte development that also recovered with age (**Figure 6**). There are numerous trophic interactions between microglia and astrocytes (Matejuk and Ransohoff, 2020) that likely contribute to this link. Importantly, in the older animals there was no evidence of astrocytosis that is commonly seen in neuroinflammation. The *Csf1r*^E631K/+^ mice are clearly different from *Csf1r*^+/-^ mice (Chitu et al., 2020; Chitu et al., 2015; Chitu et al., 2021). The *Csf1r*^ΔFIRE^ mutation, like the complete knockout, is not dosage compensated (Rojo et al., 2019). Accordingly, there is a 50% reduction in CSF1R on individual microglia. We were able to recapitulate the increased microglial density reported in the C57Bl6/J *Csf1r*^+/-^ mouse (Chitu et al., 2020; Chitu et al., 2015; Chitu et al., 2021) in heterozygous *Csf1r*^ΔFIRE^ mice on the C57Bl/6J background (**Figure S5**). We suggest the 50% loss of expression has a minimal impact on proliferation but reduces CSF1R-mediated endocytosis allowing growth factors to increase in the brain with time. Accordingly, with age a denser population of microglia can be maintained by the available CSF1 and IL34.

Neither microglia-deficient *Csf1r*^ΔFIRE/ΔFIRE^ nor CSF1-resistant *Csf1r*^E631K/+^ mice exhibit the brain pathology associated with the loss of microglia due to dominant and homozygous recessive CSF1R mutations in human patients. However, it is not clear that CSF1R mutations are strictly causal in human ALSP. There are multiple reports of asymptomatic aged individuals who carry kinase-dead mutations that are disease-associated in their progeny, siblings, or other relatives (reviewed in (Chitu et al., 2021)). CSF1R mutations might interact genetically with other common allelic variants associated with susceptibility to neurodegeneration. There is certainly evidence of epistatic interactions between *Csf1r* mutations and genetic background in mouse strains with C57BL/6J being uniquely sensitive (Chitu et al., 2016). There are obvious parallels with another adult-onset microgliopathy, Nasu-Hakola disease, associated with mutations in *TREM2* or the adaptor, *TYROBP. Trem2*^-/-^ mice do not exhibit neurodegenerative pathology but TREM2 loss-of-function sensitises to development of disease in dementia models (Filipello et al., 2018; Ulland et al., 2017)

No mouse model can recapitulate the gene-by-environment interactions that are well-documented in more common forms of neurodegenerative disease (Dunn et al., 2019) and likely also influence disease progression in ALSP. There is a massive literature on the roles of microglia in neuroprotection (Prinz et al., 2019). An inability to respond to CSF1/IL34 in the brain and/or in the periphery could predispose to brain pathology. For example, the early CSF1-dependent microglial response is essential for efficient control of Herpes simplex virus encephalitis challenge in mice (Uyar et al., 2020). Our ongoing studies address the impact of the *Csf1r*^E631K/+^ genotype in various disease models.

Notwithstanding the lack of brain pathology, the data strongly support a dominant-negative mechanism for the most common kinase-dead ALSP-associated mutations. The *Csf1r*^E631K/+^ mouse provides a novel hypomorphic model for understanding CSF1R biology and may also provide insight into the likely selective impacts of CSF1R kinase inhibitors on peripheral macrophage populations.

## Acknowledgements

The generation of the mice was funded by a grant from the Medical Research Council (MRC) UK grant MR/M019969/1 to DAH. This work was supported by Australian National Health and Medical Research Council (NHMRC) Grant GNT1163981 awarded to DAH and KMS. The laboratory receives core support from The Mater Foundation. We acknowledge input and expertise from the Biological Resources facility and the Preclinical Imaging, Microscopy and Flow Cytometry facilities of the Translational Research Institute (TRI) and the Queen’s Medical Research Institute. TRI is supported by the Australian Government. We acknowledge the Queensland Brain Institute Histology and Microscopy Facility for the use of their Stereology and Neurolucida equipment, funded by the Australian Research Council Grant LIEF LE100100074.

